# The effect of stimulation interval on plasticity following repeated blocks of intermittent theta burst stimulation

**DOI:** 10.1101/205781

**Authors:** Nga Yan Tse, Mitchell R. Goldsworthy, Michael C. Ridding, James P. Coxon, Paul B. Fitzgerald, Alex Fornito, Nigel Rogasch

## Abstract

**Introduction:** Theta burst stimulation (TBS) is a non-invasive brain stimulation paradigm capable of influencing cortical circuits in humans by inducing neural plasticity. Applying spaced blocks of TBS can affect both the direction and magnitude of plasticity, but the impact of interval duration on these interactions following intermittent TBS (iTBS) is unclear.

**Objectives:** To assess the effect of interval duration on plasticity magnitude/direction following spaced iTBS.

**Methods:** 15 healthy participants received three different iTBS conditions on separate days: single iTBS; spaced iTBS with a 5 minute interval (iTBS-5); and spaced iTBS with a 15 minute interval (iTBS-15). Changes in cortical excitability and short-interval cortical inhibition (SICI) resulting from iTBS were assessed via motor-evoked potentials (MEPs) measured from the first dorsal interosseus muscle before and up to 60 mins following stimulation.

**Results:** iTBS-15 increased MEP amplitude up to 60 mins post stimulation, whereas iTBS-5 decreased MEP amplitude. In contrast, MEP amplitude was not altered by single iTBS. Despite the significant effect of iTBS-15 on MEP amplitude at the group level, there was still considerable inter-individual variability, with only 53% of individuals meeting response criteria. Modulation of SICI did not differ between conditions.

**Conclusions:** The interval duration between spaced iTBS plays an important role in determining the direction of plasticity on excitatory, but not inhibitory circuits in human motor cortex. While iTBS-15 can increase the magnitude of facilitation in some individuals compared to single iTBS, this approach still suffers from high inter-individual variability.

## INTRODUCTION

The capacity of the brain to modulate the strength of synaptic connections, commonly called synaptic plasticity, is a fundamental mechanism for healthy brain functioning, representing a key neural substrate for learning, memory, and development [1]. Changes in synaptic strength are governed by a variety of mechanisms, of which long-term potentiation (LTP) and depression (LTD) regulated by voltage-dependent n-methyl-d-aspartate (NMDA) receptors are the best characterised [2]. In *in vitro* studies of neural tissue, LTP is observed naturally following learning [3], but can also be experimentally induced by external stimulation delivered at certain patterns mimicking natural brain rhythms. For example, theta-burst stimulation (TBS), which is a widely used plasticity-inducing paradigm, involves nesting high frequency (e.g. 100 Hz) stimulation pulses within slower frequencies (e.g. 5 Hz) [4,5]. However, the direction, magnitude, and duration of plasticity induced either naturally or experimentally is highly dependent on the history of synaptic activity, a phenomenon known as metaplasticity [6]. For instance, if a TBS paradigm which results in LTP is immediately followed by a second round of TBS, further LTP induction is blocked or even reversed [7,8]. Such interactions represent a homeostatic process that are thought to regulate the induction of LTP, preventing run-away changes in cortical excitability which could lead to excitotoxicity [6]. However, if the interval between the TBS protocols is increased, a longer more stable form of LTP termed “late-LTP” is induced [7]. This non-homeostatic form of metaplasticity is likely important for the persistent changes in synaptic strength required for learning and memory storage [9]. Thus, the timing of the interval between stimulation trains plays a critical role in determining the type of plasticity that can be induced [7,10].

Plasticity can be studied in humans using non-invasive brain stimulation methods, such as transcranial magnetic stimulation (TMS) [11]. TBS protocols delivered with TMS (typically 50 Hz bursts at 5 Hz for 600 pulses) can alter cortical excitability beyond the period of stimulation and are blocked by NMDA antagonists [12], reminiscent of LTP/LTD-like plasticity observed in animal experiments [13]. Early reports suggested that an intermittent pattern of stimulation (2 s on, 8 s off; iTBS) resulted in increased cortical excitability similar to LTP, whereas a continuous pattern (no off period) resulted in decreased excitability similar to LTD [14]. However more recent studies have reported considerable inter-individual variation in response direction [15–18]. When blocks of TBS are repeated in humans, evidence for both homeostatic (e.g. plasticity blocked or direction reversed), and non-homeostatic (e.g. increased magnitude) interactions have been reported [19], however the factors which govern the direction and magnitude of metaplasticity in humans are unclear. Surprisingly, few studies have systematically assessed the impact of altering the interval between repeated TBS blocks on plasticity in humans [20]. Further, it is unclear whether inhibitory circuits, which are also likely targeted by TBS [21], show metaplasticity. Given the interest in using iTBS as a clinical tool to treat disorders such as depression [22], which would benefit from the reliable and stable induction of plasticity conferred by late-LTP, it is important to understand how such parameter choices impact the outcome of stimulation.

The aims of this study were twofold. First, we assessed whether the interval between consecutive spaced iTBS blocks affects the direction and magnitude of changes in cortical excitability following stimulation. Second, we assessed whether plasticity on the inhibitory circuits targeted by iTBS is also altered following spaced stimulation. Given the findings from animal studies [e.g. 7], we hypothesised that shorter intervals (5 mins) would lead to reductions in MEP amplitude and SICI, whereas longer intervals (15 mins) would result in increased MEP and SICI facilitation compared to a single block of iTBS.

## MATERIALS AND METHODS

### Participants

20 healthy participants were recruited for this study. Prior to the experiment, all participants were screened for any contraindications to TMS based on the TMS safety guidelines [23], and provided written informed consent. Five participants were withdrawn from the study due to high stimulation thresholds (n=2), inability to complete all sessions (n=1), light-headedness following resting motor threshold (n=1), and persistent muscle activity during TBS (n=1). As such, 15 participants completed all three experimental conditions (mean age = 24.8 ± 4 years; 7 females; 1 left handed). The study was approved by the Monash University Human Research Ethics Committee.

### Electromyography

Participants were seated comfortably in a chair with their right hand and arm resting on a pillow. Surface electromyography (EMG) was recorded from the right first dorsal interosseous (FDI) muscle. Ag-AgCl electrode pairs were placed in a belly-tendon montage over the FDI muscle, with a ground electrode placed over the distal surface of the hand. EMG signals were recorded from 100 ms before to 500 ms after each TMS pulse. Signals were amplified (×1000), bandpass filtered (10-1000 Hz), digitized (5 kHz), and stored on a computer for offline analysis (Powerlab 26T, ADInstruments Ltd., New Zealand).

### Transcranial magnetic stimulation

TMS was applied through a figure-of-eight coil (C-B60; 75mm outer diameter) connected to a MagPro X100 stimulator (MagVenture, Denmark). The current waveform was biphasic for all conditions. Biphasic pulses were chosen over monophasic pulses to more accurately index cortical excitability from the neural pool stimulated using TBS (which uses biphasic pulses). The coil was held tangentially to the skull with the handle pointing backwards, at an angle of 45° to the sagittal plane, such that an anterior-posterior followed by a posterior-anterior current flow was induced in the underlying cortex. The coil was held over the hand area of the left motor cortex. The scalp position resulting in the most consistent and largest MEP in the FDI muscle (i.e. the motor ‘hotspot’) was determined and used throughout the session. This position was marked on the participant’s scalp with a water-soluble pen in order to keep coil positioning consistent. The resting motor threshold (RMT) was then determined as the minimum intensity necessary to elicit at least 5 of 10 MEPs with a peak-to-peak amplitude >50 μV while the target muscle is relaxed. The stimulus intensity was then increased to induce an MEP of ~1 mV in amplitude (stimulus intensity; S1mV). This stimulus intensity was used throughout the experiment to index changes in cortical excitability.

Intracortical inhibition was assessed using the paired-pulse short-interval cortical inhibition (SICI) paradigm. SICI was applied with an inter-stimulus interval of 2 ms. This interval was chosen to index true GABA_A_-mediated neurotransmission and limit contamination from short-interval cortical facilitation [24]. The conditioning pulse intensity was set at 70% of RMT, and the test pulse intensity at S1mV. The conditioning intensity was chosen to match the intensity used for iTBS (see below), thereby allowing a direct evaluation of excitability changes induced in the neural population targeted by TBS. As the amplitude of the test MEP can influence SICI magnitude [25], SICI recordings were repeated in the post iTBS blocks with the test pulse stimulus adjusted to provide an MEP of ~1 mV in amplitude to a single TMS pulse (MEP_adj_ and SICI_adj_). MEPs and SICI were assessed at baseline, and at 5, 15, 30, 45, and 60 mins post the main iTBS session. 15 trials were collected for each measurement at a frequency of 0.2 Hz.

### Intermittent theta burst stimulation

iTBS was delivered over the FDI motor hotspot using an actively cooled figure-of-eight magnetic coil (MCF-B65, MagVenture; 75mm outer diameter). RMT was re-measured using the cooled coil. Stimulation consisted of a burst of three pulses administered at 50 Hz, repeated at a frequency of 5 Hz, delivered in 2 s trains followed by an 8 s interval for a total of 600 pulses [14]. Stimulation intensity was set at 70% of RMT. The more common TBS intensity of 80% active motor threshold was not used to avoid voluntary contractions prior to TBS, as several studies have suggested that such contractions can interfere with subsequent TBS-induced plasticity [26–28].

To address the primary and secondary aim of the study, changes in cortical excitability and inhibition were assessed after three different iTBS conditions: a single iTBS session (iTBS); iTBS primed by another iTBS session 5 minutes prior (iTBS-5); and iTBS primed by another iTBS session 15 minutes prior (iTBS-15). To ensure each condition was as similar as possible, sham iTBS was applied at the 5 and 15 min intervals in the conditions not requiring active stimulation at these time points (figure 1). Sham iTBS was achieved by rotating the coil head 90° around its axis so the magnetic field ran perpendicular to the scalp, and the coil wing rested over the motor hotspot. The three conditions were randomised and completed on different days separated by at least 72 hours.

**Figure 1:**
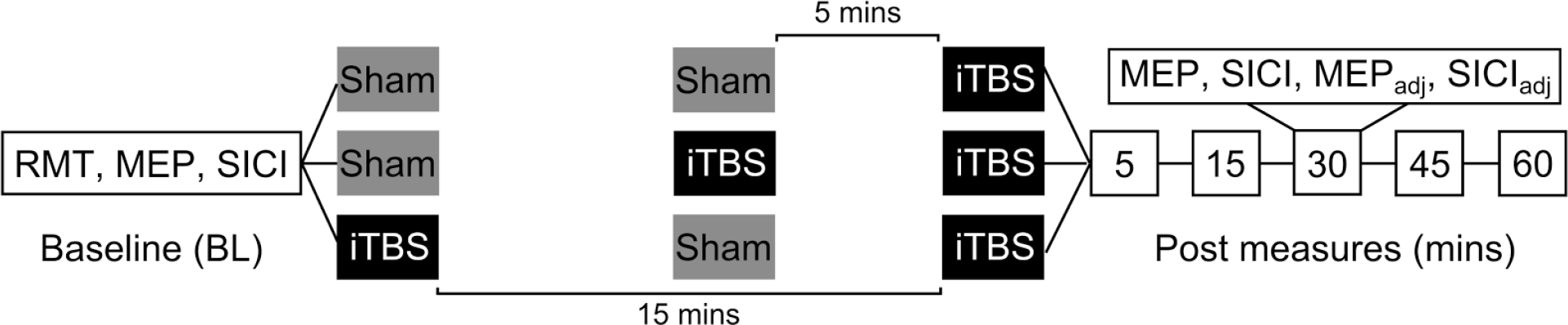
Diagram demonstrating the study protocol. Participants received three different iTBS conditions on separate days: single iTBS (top); iTBS-5 (middle); and iTBS-15 (bottom). Note that time is not drawn to scale for baseline and post measures.

### Data analysis

Trials with background muscle activity (root-mean-squared EMG >10 μV in the 100 ms prior to stimulation) were excluded from further analysis. To index changes in cortical excitability following iTBS, peak-to-peak MEP amplitudes were averaged across trials in each recording block following single pulse TMS. To compare changes in cortical excitability between each condition, post MEP amplitudes were normalised to baseline, and averaged across post iTBS time period for each individual (grand average response). SICI was quantified by dividing the mean conditioned peak-to-peak MEP amplitude (e.g. the test MEP following paired-pulse TMS), by the mean peak-to-peak amplitude of the single pulse MEP alone. For the post iTBS blocks, this was repeated for trials using the adjusted test MEP amplitude to calculate SICI_adj_. Using this formula, values close to 0 indicate strong inhibition, whereas values close to 1 indicate weak inhibition.

### Statistics

One-way analysis of variance (ANOVA) was used to compare baseline RMT, S1mV, MEP amplitude and SICI between conditions. To examine the effect of different spaced iTBS intervals on cortical excitability and inhibition, 3 × 6 repeated measures ANOVAs (RMANOVAs) were used to test the main effects of CONDITION (iTBS, iTBS-5, iTBS-15) and TIME (baseline, 5, 15, 30, 45, 60) on MEP amplitude, SICI, MEP_adj_ amplitude and SICI_adj_. The Shapiro-Wilk test was used to assess normality and detected outliers in non-normal data were winsorised to the next highest value. Mauchly’s test was used to assess sphericity and the Greenhouse-Geiser correction applied if necessary. In the case of significant main effects or interactions, targeted post-hoc testing was applied using Fisher’s PLSD test. Pearson’s correlations were used to assess whether there was any relationship between grand average changes in MEP amplitude between conditions. In all tests, a value of p ≤ 0.05 was considered statistically significant. All data are expressed as mean ± SD in the text and mean ± SEM in the figures.

## RESULTS

### Baseline measures

There were no differences in RMT (F_2,14_=0.3, p=0.736), S1mV intensity (F_2,14_=0.3, p=0.764), MEP amplitude (F_2,14_=1.9, p=0.174), or SICI (F_2,14_=1.4, p=0.246) between conditions at baseline.

### Cortical excitation following single and spaced iTBS

To assess whether the interval between repeated blocks of iTBS influences the direction of plasticity, we first compared iTBS-induced changes in MEP amplitude between conditions. RMANOVA on MEP amplitude revealed a significant main effect of CONDITION (F_2,14_=7.4, p=0.003, η^2^=0.35), no effect of TIME (F_5,14_=0.3, p=0.933, η^2^=0.02), but most importantly, a significant CONDITION × TIME interaction (F_10,14_=2.4, p=0.011, η^2^=0.15). For the main effect of CONDITION, post-hoc tests showed that MEP amplitudes were larger following iTBS-15 than both single iTBS (p=0.042) and iTBS-5 (p=0.004). For the interaction, post-hoc tests showed that MEP amplitudes were larger following iTBS-15 compared to iTBS-5 at 30 mins (p=0.001), 45 mins (p=0.045) and 60 mins (p=0.002), and were larger following iTBS-15 (p=0.034) and smaller following iTBS-5 (p=0.012) compared to single iTBS at 60 mins (figure 2). Secondary analyses confirmed that, compared to baseline, MEP amplitudes were increased at 30 mins (p=0.033) and 60 mins (p=0.049) following iTBS-15, and decreased at 30 mins (p=0.030) and 60 mins (p<0.001) following iTBS-5. No changes were identified following single iTBS (all p>0.05). These results suggest that the spacing interval between repeated applications of iTBS effects the direction of excitability change, particularly at later time points.

**Figure 2:**
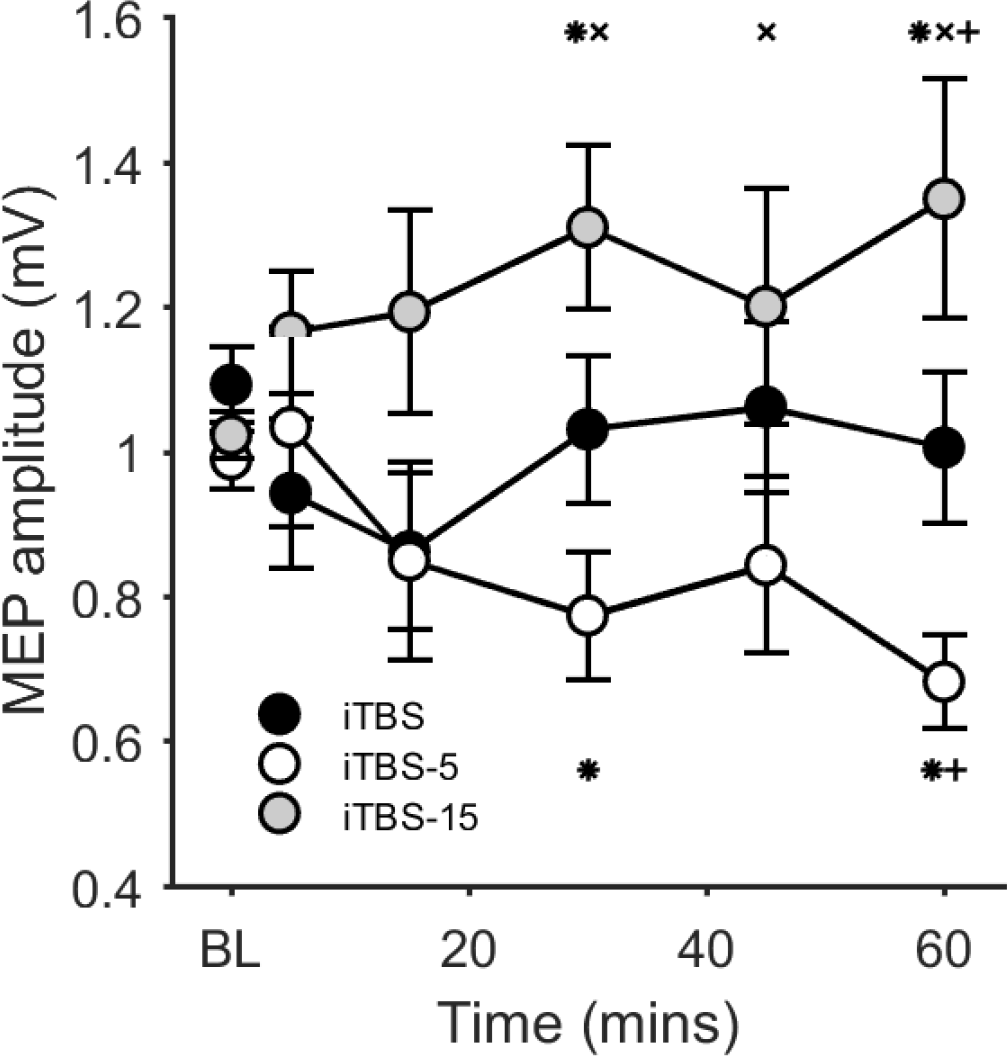
MEP amplitudes following single and spaced iTBS. BL = baseline. * p<0.05 compared to BL; × p < 0.05 compared to iTBS-5; + p < 0.05 compared to single iTBS.

### Individual response to spaced iTBS

Figure 3 shows individual responses to iTBS for each condition normalised to baseline. Considerable inter-individual variability is apparent in response to both single and spaced iTBS conditions (figure 3A-C). To quantify the percentage of ‘responders’, ‘non-responders’, and ‘opposite responders’ to each condition, normalised changes in MEP amplitude were averaged across post iTBS time points to generate a grand average value for each individual. As we expected an increase in MEP amplitude to iTBS, individuals with a grand average value >1.1 were considered responders, <0.9 opposite responders, and 0.9<grand average<1.1 as non-responders for the single iTBS and iTBS-15 conditions according to a previous definition [18,29]. Given that we found that the iTBS-5 condition reduced excitability at the group level, we instead considered individuals with a grand average value <0.9 (i.e. a reduction in MEP amplitude) as responders to this condition, and >1.1 opposite responders. Only 33% of individuals were deemed responders to single iTBS (figure 3D). Although iTBS-15 resulted in a larger increase in MEP amplitude at the group level, the number of responders at the individual level was only marginally higher than single iTBS, with 53% of individuals deemed responders (figure 3F). Similar to the other conditions, only 40% of individuals were deemed responders to iTBS-5 with reductions in MEP amplitude (figure 3E).

**Figure 3:**
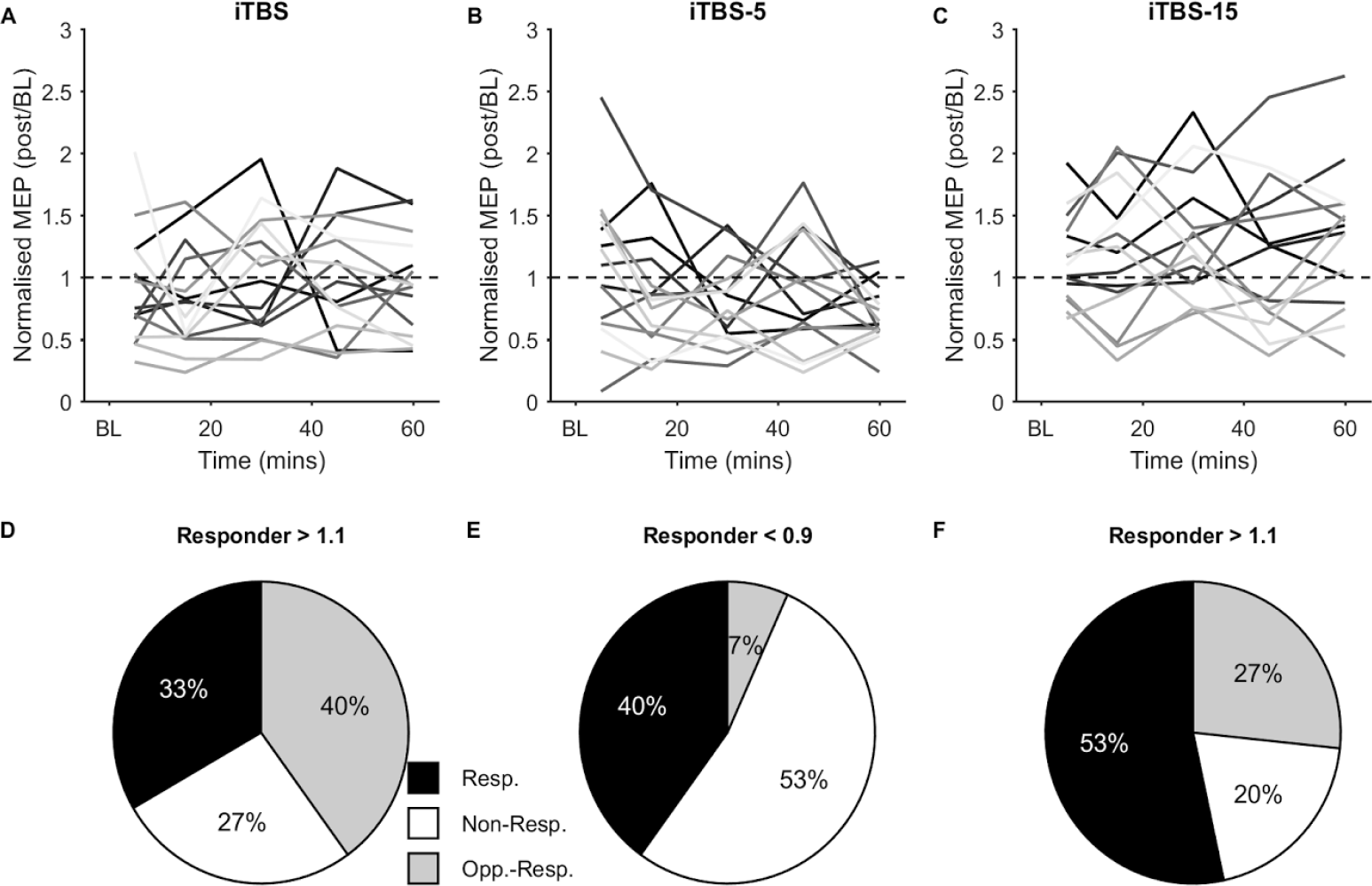
Individual responses to single and spaced iTBS. A-C: Normalised changes in MEP amplitude following different iTBS conditions in individuals, represented as different shades of grey. Changes in MEP amplitude have been normalised to baseline values, with score >1 indicating an increase, and <1 a decrease in excitability. D-F: Percentage of ‘responders (resp.)’, ‘non-responders (non-resp.)’, and ‘opposite responders (opp.-resp.)’ to each iTBS condition. Responders are defined as having a grand average response (average of all normalised post iTBS MEP amplitudes) >1.1 for iTBS and iTBS-15 and <0.9 for iTBS-5, whereas opposite responders have a response <0.9 for iTBS and iTBS-15 and >1.1 for iTBS-5.

### Correlations between changes in MEP amplitude following spaced iTBS

To assess whether responders and non-responders were similar between different iTBS conditions, we correlated the grand average response between single and spaced iTBS. There was no relationship between the responses to different conditions for any pairing (iTBS v iTBS-5, p=0.76; iTBS v iTBS-15, p=0.56; iTBS-5 v iTBS-15, p=0.78; figure 4), suggesting that different mechanisms may underlie the variability in response to single and spaced iTBS.

**Figure 4:**
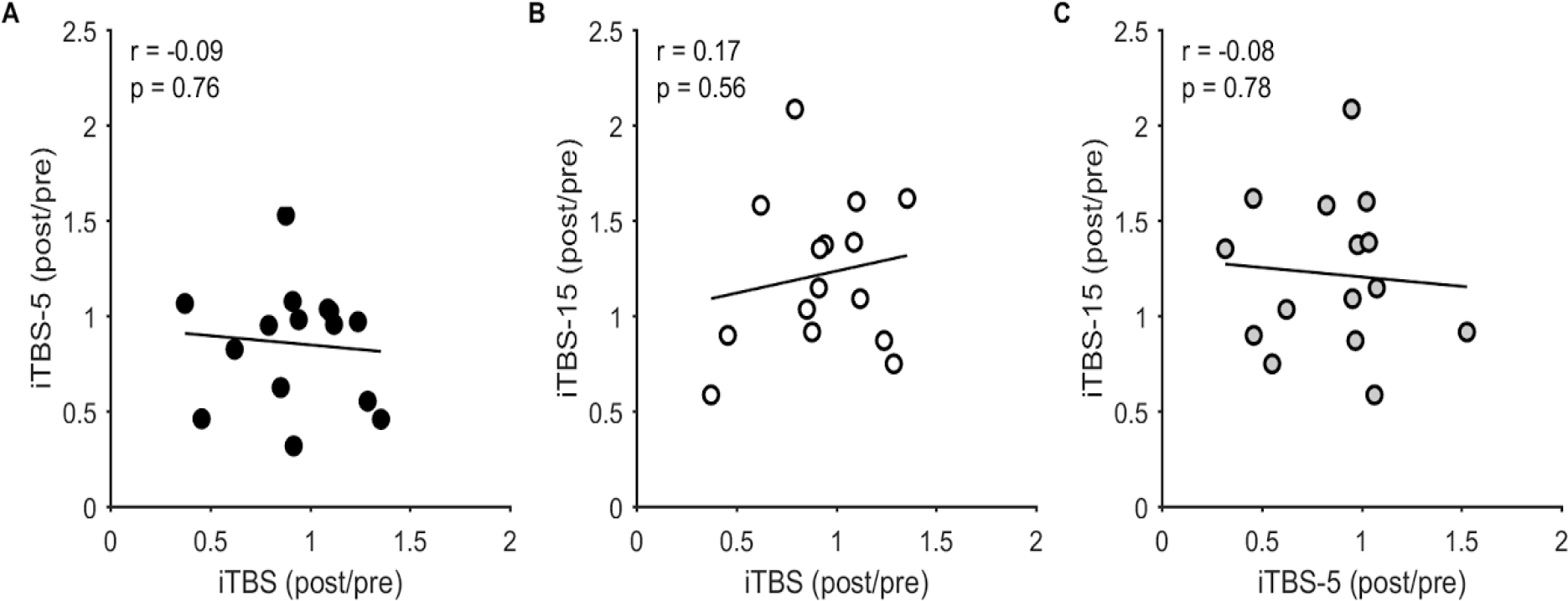
Relationship between single and spaced iTBS. Correlations between the normalised grand average MEP amplitude following different iTBS conditions.

### Cortical inhibition following single and spaced iTBS

To assess whether similar interactions between plasticity and interval duration occur within inhibitory populations, we also assessed changes in SICI following single and spaced iTBS. RMANOVA on SICI revealed no main effect of CONDITION (F_2,14_=3.3, p=0.052, η^2^=0.19) or TIME (F_5,14_=1.5, p=0.206, η^2^=0.10), and no CONDITION × TIME interaction (F_10,14_=1.3, p=0.274, η^2^=0.09) (figure 5A). To control for possible confounding effects of iTBS-induced changes in MEP amplitude on SICI, we repeated SICI measurements (SICI_adj_) with MEP amplitude adjusted to 1 mV (MEP_adj_). This adjustment to the test stimulus intensity successfully matched the non-conditioned response amplitude as evidenced by no main effect of CONDITION (F_2,14_=1.9, p=0.164, η^2^=0.12) and no CONDITION × TIME interaction (F_10,14_=0.4, p=0.944, η^2^=0.02) on adjusted MEP amplitude following iTBS (figure 5B). As with unadjusted SICI, there was no main effect of CONDITION (F_2,14_=1.2, p=0.238, η^2^=0.09), or CONDITION × TIME interaction (F_10,14_=3.3, p=0.274, η^2^=0.09) on SICI_adj_, however there was a main effect of TIME (F_5,14_=1.5, p=0.042, η^2^=0.19). Averaged across conditions, SICI_adj_ increased at 5 mins (p=0.009), 30 mins (p=0.034), 45 mins (p=0.019), and 60 mins (p=0.004) following iTBS (figure 5C). These findings suggest that inhibitory circuits do not show similar changes in plasticity following spaced iTBS at different intervals as excitatory populations.

**Figure 5:**
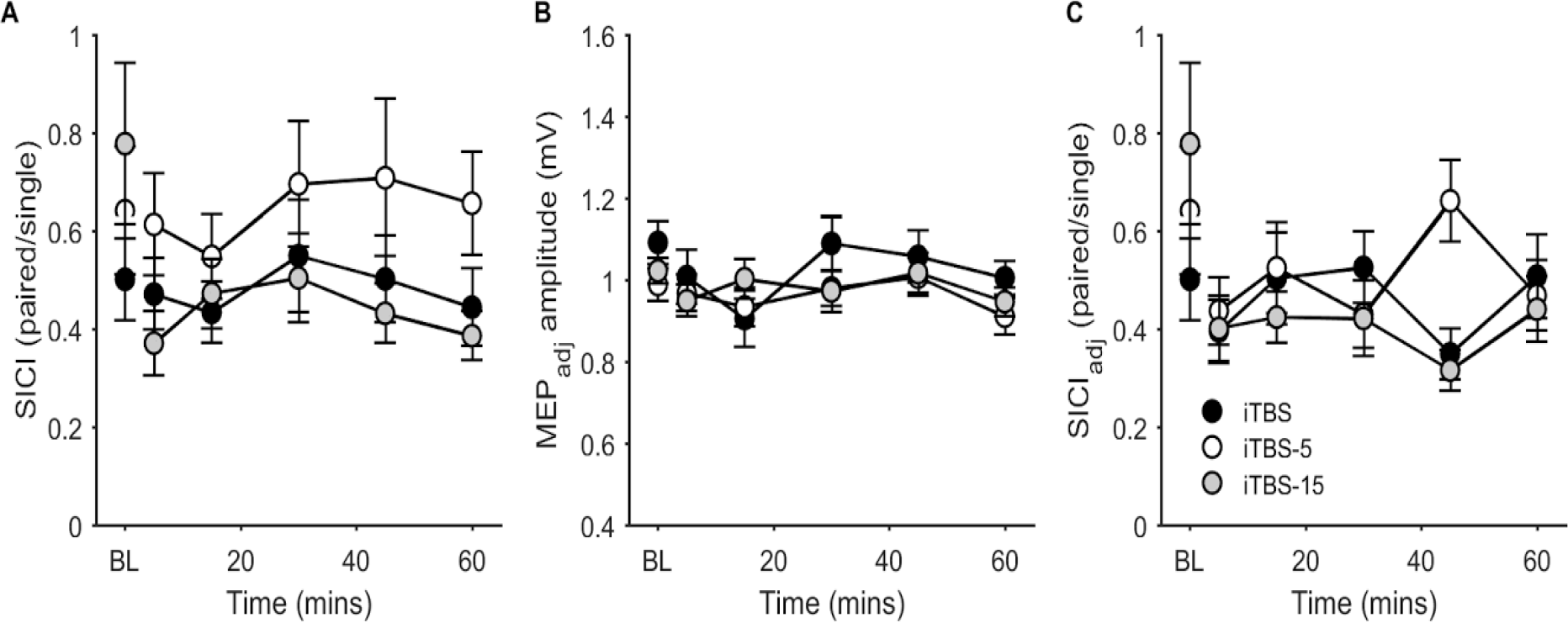
SICI following single and spaced iTBS. A) Changes in SICI following iTBS without adjusting test MEP amplitude. B) MEP amplitudes at each time point after adjusting TMS intensity to give a 1 mv response (MEP_adj_). C) Changes in SICI following iTBS after adjusting test MEP amplitude (SICI_adj_).

Although there was no significant difference in baseline SICI between conditions, inspection of figure 5 suggested that baseline SICI was qualitatively lower (i.e. closer to 1) in the iTBS-15 condition compared to single iTBS. To ensure that lower SICI was not driving the increase in MEP amplitude in the iTBS-15 condition, we correlated SICI strength at baseline with grand average change in normalised MEP amplitude following iTBS. There was no significant correlation between baseline SICI and normalised changes in MEP amplitude for combined single and iTBS-15 conditions (r=0.12, p=0.54) or iTBS-15 alone (r=0.07, p=0.81), suggesting between session differences in baseline SICI were not driving the larger increases in MEP amplitude following iTBS-15.

## DISCUSSION

Repeating blocks of TBS can influence the magnitude and direction of plasticity in humans, however the factors determining the nature of this interaction are not well defined. In this study, we have found that the interval duration between repeated blocks of iTBS impacts the resulting plasticity direction, with shorter intervals (5 mins) reducing cortical excitability, and longer intervals (15 mins) increasing excitability. Unlike cortical excitability, we could not find any evidence that plasticity of inhibitory circuits is altered following repeated blocks of iTBS. Despite larger MEP facilitation at the group level following repeated iTBS with a 15 minute interval compared to single iTBS, only approximately half of the sample could be considered ‘responders’, indicating that spaced iTBS still suffers from large inter-individual variability.

### Cortical excitability following single iTBS

A single block of iTBS (600 pulses) is reported to increase cortical excitability indexed by MEP amplitude for ~30 mins following stimulation [14], changes which are blocked by NMDA receptor antagonists [12]. As such, cortical excitability increases following iTBS are thought to reflect LTP-like plasticity mechanisms [30]. We could not find any evidence for changes in cortical excitability following a single block of iTBS at the group level. Indeed, only 33% of individuals in our study could be considered responders, with 40% responding in the opposite direction. This finding may seem contradictory to the expected facilitatory effect of iTBS [14], however several recent studies with large sample sizes (N>50) have reported considerable inter-individual variability in response to iTBS, resulting in no net differences in excitability at the group level [15,16]. A recent meta-analysis found evidence for publication bias in the iTBS literature [31], suggesting inter-individual variability in response to a single block of iTBS may be larger than initially expected. Our findings are consistent with a growing body of literature suggesting large inter-individual variability in response to a single block of iTBS.

### The duration between repeated blocks of iTBS impacts plasticity direction and magnitude

In animal slice studies, the instability/variability of plasticity induced by a single block of stimulation is overcome by administering multiple blocks of stimulation to induce late-LTP [10], in which the interval duration plays an important role in determining the magnitude and direction of plasticity [7]. In humans, two blocks of cTBS at 10-15 minute intervals results in a more reliable and longer lasting reduction in MEP amplitude [32] and impairment in behaviour [33] compared with a single block alone (however see [34]). Importantly, the plasticity following spaced cTBS is resistant to reversal by subsequent physiological activity, such as muscle contractions, suggesting consolidation of LTD-like plasticity [35]. In contrast, reducing the interval duration to 2 or 5 minutes [20], or increasing the interval to 25 minutes blocks LTD-like plasticity following spaced cTBS [36] consistent with homeostatic plasticity.

The interactions between repeated blocks of iTBS on plasticity is less clear. Previous studies report that plasticity is blocked following 5 minute interval spaced iTBS [20], and facilitated/prolonged with either 10 minute intervals [37] or 3 blocks of 15 minute intervals [38]. Furthermore, repeated blocks of facilitatory paired associative stimulation, another TMS paradigm which induces LTP-like plasticity, prolongs plasticity duration [39]. In contrast, several studies have reported blocking or reversal of plasticity following repeated iTBS intervals at 15 [34], 20 [20] and 25 minutes [36]. An important difference between studies reporting homeostatic and non-homeostatic interactions at longer iTBS intervals is the performance of a voluntary contraction at baseline during motor threshold determination (homeostatic; [20,34,36]) compared to remaining at rest (non-homeostatic; our study, [37,38]). Performing muscle contractions prior to TBS can influence subsequent plasticity induction [26–28,32], therefore complicating the interpretation of changes in excitability. As such, we designed our study to avoid the voluntary muscle activity associated with determination of active motor threshold. We have shown that, unlike a single block of iTBS, two blocks of iTBS separated by a 15 minute interval increased MEP amplitudes at the group level, which persisted for 60 minutes following stimulation. In contrast, shortening the interval to 5 minutes reversed the direction of plasticity, instead reducing MEP amplitude. Our findings suggest that the interval between repeated blocks of iTBS is important for determining the direction of subsequent plasticity.

Another motivation for exploring spaced iTBS is to assess whether this method improves the response rate by reducing inter-individual variability compared to single iTBS. Such improvements are essential for the development of TBS as a potential clinical tool for the treatment of brain disorders such as depression [22]. Indeed, one study found that spaced cTBS resulted in 100% of participants showing reduced MEP amplitude [32] compared to 58% with single cTBS. We found that 53% of participants had the desired facilitatory response following iTBS-15. While this was a marginal increase on the response rate to single iTBS (33%) in our sample, the inter-individual variability to iTBS-15 is still high. A similar result was reported following three blocks of iTBS-15, which showed that, while spaced iTBS increased the magnitude of the response, it did not convert non-responders to responders [29]. Taken together, our results suggest that while iTBS-15 can increase the magnitude and duration of plasticity effects in some individuals, this method still suffers from high inter-individual variability, which may limit its clinical potential.

### Plasticity of inhibitory circuits following repeated iTBS

In addition to altering cortical excitability, there is also some evidence that iTBS can alter inhibitory circuits. Huang and colleagues reported that a single block of iTBS increased GABA_A_-mediated neurotransmission as assessed using SICI [14], however, a recent meta-analysis found no evidence for changes in SICI following iTBS across 13 datasets [31]. We also found no evidence for changes in SICI following either single or spaced iTBS. However we did find a general increase in SICI when we adjusted the test MEP amplitude to account for iTBS-induced changes in cortical excitability, which did not differ between conditions. Goldsworthy and colleagues also found that single and spaced cTBS had similar effects on inhibitory circuits, with both conditions decreasing SICI [40]. In contrast, Murakami and colleagues found that spaced iTBS with a 15 minute interval resulted in reduced SICI compared to a single block of iTBS [34]. The reasons for this discrepancy are unclear, however the inclusion of voluntary contractions prior to TBS may have influenced the outcomes of the latter study. We also assessed whether baseline SICI could explain the facilitation in MEPs following iTBS-15, as lower inhibition can influence subsequent plasticity. In line with other studies using single iTBS [16], baseline SICI did not predict response to iTBS-15. Taken together, our results suggest that spaced iTBS does not result in metaplasticity of inhibitory circuits, at least at the intervals tested.

### Limitations

There are several limitations to the current study. First, we did not assess changes in MEP amplitude or SICI directly following the priming block of stimulation, as we did not want to inadvertently disrupt any ongoing plasticity. Although results from the single iTBS condition suggest that MEP amplitude and SICI were unaffected by one block of stimulation in our sample, we are unable to definitely assess whether priming stimulation did or did not alter cortical excitability or inhibition. Importantly, metaplasticity is not dependent on changes in synaptic efficacy following priming [6], and several studies in human motor cortex have reported metaplasticity following repeated TBS blocks without excitability changes following the priming block [32,34,37]. Second, we used biphasic TMS pulses to assess MEP amplitude and SICI following iTBS, whereas the majority of other studies have used monophasic pulses for these assessments. The choice to use biphasic pulses was deliberate, as we wanted to more accurately assess the neural populations targeted by iTBS, which are stimulated using biphasic pulses. Changes in MEP amplitude following TBS assessed with biphasic pulses are highly comparable to those using monophasic pulses [36]. As such, it is unlikely this choice had a major impact on the study outcomes and conclusions.

## CONCLUSIONS

We provide evidence that the interval duration between repeated blocks of iTBS over human motor cortex determines the direction of plasticity on excitatory circuits, with shorter intervals favouring reduced excitability, and longer intervals facilitated excitability. In contrast, repeated blocks of iTBS did not result alter plasticity of inhibitory circuits. Although spaced iTBS with a 15 minute interval increases the magnitude of MEP facilitation at the group level, substantial inter-individual variability still exists, with just over half of the population responding in the desired direction to stimulation. Understanding the determinants of this variability in response to iTBS will be essential for designing stimulation paradigms to more consistently drive plasticity in a given individual.

## ACKNOWLEDGMENTS

This work was supported by the National Health and Medical Research Council (NHMRC) of Australia via a Project Grant awarded to AF and NCR (1104580). MRG was supported by an NHMRC-Australian Research Council (ARC) Dementia Research Development Fellowship (1102272), AF an ARC Fellowship (FT130100589) and NCR by an NHMRC Early Career Fellowship (1072057).

